# Strength in numbers: Large-scale integration of single-cell transcriptomic data reveals rare, transient muscle progenitor cell states in muscle regeneration

**DOI:** 10.1101/2020.12.01.407460

**Authors:** David W. McKellar, Lauren D. Walter, Leo T. Song, Madhav Mantri, Michael F.Z. Wang, Iwijn De Vlaminck, Benjamin D. Cosgrove

## Abstract

Skeletal muscle repair is driven by the coordinated self-renewal and fusion of myogenic stem and progenitor cells. Single-cell gene expression analyses of myogenesis have been hampered by the poor sampling of rare and transient cell states that are critical for muscle repair, and do not provide spatial information that is needed to understand the context in which myogenic differentiation occurs. Here, we demonstrate how large-scale integration of new and public single-cell and spatial transcriptomic data can overcome these limitations. We created a large-scale single-cell transcriptomic dataset of mouse skeletal muscle by integration, consensus annotation, and analysis of 23 newly collected scRNAseq datasets and 79 public single-cell (scRNAseq) and single-nucleus (snRNAseq) RNA-sequencing datasets. The resulting compendium includes nearly 350,000 cells and spans a wide range of ages, injury, and repair conditions. Combined, these data enabled identification of the predominant cell types in skeletal muscle with robust, consensus gene expression profiles, and resolved cell subtypes, including endothelial subtypes distinguished by vessel-type of origin, fibro/adipogenic progenitors marked by stem potential, and many distinct immune populations. The representation of different experimental conditions and the depth of transcriptome coverage enabled robust profiling of sparsely expressed genes. We built a densely sampled transcriptomic model of myogenesis, from stem-cell quiescence to myofiber maturation and identified rare, short-lived transitional states of progenitor commitment and fusion that are poorly represented in individual datasets. We performed spatial RNA sequencing of mouse muscle at three time points after injury and used the integrated dataset as a reference to achieve a high-resolution, local deconvolution of cell subtypes. This analysis identified the temporal variation in the colocalization of immune cell subtype interactions with myogenic progenitors during injury recovery. We provide a public web tool to enable interactive exploration and visualization of this rich single-cell transcriptomic resource. Our work supports the utility of large-scale integration of single-cell transcriptomic data as a tool for biological discovery.

## INTRODUCTION

Muscle stem cells (MuSCs) are essential for muscle homeostasis and repair. MuSCs are typically quiescent in homeostasis and are activated in muscle damage. Their subsequent proliferation, differentiation, commitment, and fusion replenishes skeletal muscle tissue in a complex, coordinated process^1–3^. MuSCs are a rare cell type, accounting for less than 1% of the cells within skeletal muscle at homeostasis. Even rarer are the short-lived progenitor cell states between quiescent MuSCs and fully differentiated myofiber cells. Consequently, MuSCs and muscle progenitor cells (comprised of myoblasts and myocytes) are difficult to study in their native tissue context. Conventional strategies to study MuSCs and muscle progenitor cells rely on enrichment by fluorescence-activated cell sorting using transgenic reporter or prospective isolation markers^4^. These methods however are ill-suited to capture the subtle, continuous cell state transitions which are critical for myogenesis due to a paucity of highly stage-specific cell isolation markers and the rarity of these cells.

Single-cell RNA sequencing (scRNAseq) enables a detailed characterization of cell types and states in complex tissues without the need for targeted cell enrichment^5–8^. Skeletal muscle has been the focus of a number of recent scRNAseq studies, which have aimed to catalog its dynamic and heterogeneous constituent cell types and the progression of myogenic stem and progenitor cell regulation in muscle development and repair^7^. Single-nucleus RNA sequencing (snRNAseq) has been used to capture transcriptomic signatures from mature myofiber nuclei, which are largely lost during cell isolation required for scRNAseq^9–13^. Yet, despite advances in the scale of sc/snRNAseq technologies (10^3^-10^4^ cells per experiment), these methods still poorly sample rare cell types and transient cell states in detail without purification steps which can introduce marker bias and technical artefacts^14^. For example, we previously used scRNAseq to study the dynamics of hindlimb skeletal muscle regeneration in adult mice and resolved ∼12 muscle-resident cell types from ∼35,000 single-cell transcriptomes^15^. However, we observed fewer than 100 committed and fusing myogenic cells even though we sampled critical time-points of myogenic differentiation post-injury^15^. Similar studies likewise reported an infrequent sampling of committed myogenic progenitors from whole muscle samples^15–17^.

To overcome these challenges, we sought to integrate and evaluate a more diverse collection of single-cell transcriptomic data. We collected ∼95,000 additional single-cell transcriptomes from 23 new samples of regenerating mouse hindlimb muscles in older mice. We then leveraged recent improvements in batch-correction algorithms^18,19^ to incorporate 79 publicly available sc/snRNAseq datasets from 14 prior studies in published reports and public repositories in our analysis^15–17,20–28^ (**Table S1**). This led to a data compendium that included ∼350,000 cells/nuclei after quality filtering and allowed us to study the cellular composition and dynamics in response to skeletal muscle injury as function of a wide range of experimental conditions. The high depth of transcriptome coverage achieved by large-scale integration of single-cell transcriptomic data enabled us to robustly characterize rare, short-lived cell states on the myogenic cell differentiation trajectory. We identified transcription factors and surface markers that distinguish committed myoblasts (∼18 per sample, on average) and fusing myocytes (∼8 per sample, on average), which represent only 0.5% and 0.2% of all cells in the integrated muscle compendium, respectively. We performed spatial RNA sequencing of mouse muscle at three time points after injury and used the integrated compendium as a reference to achieve a high-resolution, local deconvolution of cell subtypes. Our analysis also brings new insights into the dynamics of stromal and immune cell colocalization with transient myogenic cell states.

## RESULTS

### Large-scale integration enables a high-resolution view of skeletal muscle

To profile skeletal muscle homeostasis and repair more deeply, we performed single-cell RNA sequencing on 23 adult mouse skeletal muscle samples using the 10x Chromium v3 platform. In addition to uninjured controls, we induced muscle damage in adult (7 mo) and aged (20 mo) C57BL/6J mice using notexin and collected tibialis anterior muscles at several time points within one week after injury (sample details in **Table S1, Sup. File 1**). To augment these data, we curated 79 publicly available mouse skeletal muscle sc/snRNAseq datasets that were generated on the 10x Chromium platform (v2 or v3) from PanglaoDB^29^ and SRA as of July 31, 2020. Herein we refer to each dataset by its citation, using the first name(s) listed in either the publication or deposition (for data not published) and year of release. Together, these comprised 102 individual samples with a total of 469,603 cell barcodes, before quality control and filtering (**Figs. 1A and S1, Table S1**). These data vary across sex, age (10 days to 30 months of age), chemical injury model (notexin and cardiotoxin), injury-response timepoint (0.5 to 21 days post-injury [dpi]), and sample preparation strategy, including whole-muscle dissociations and FACS enrichment of specific cell types (**Figs. 1B and Fig. S1**).

**Figure 1.**
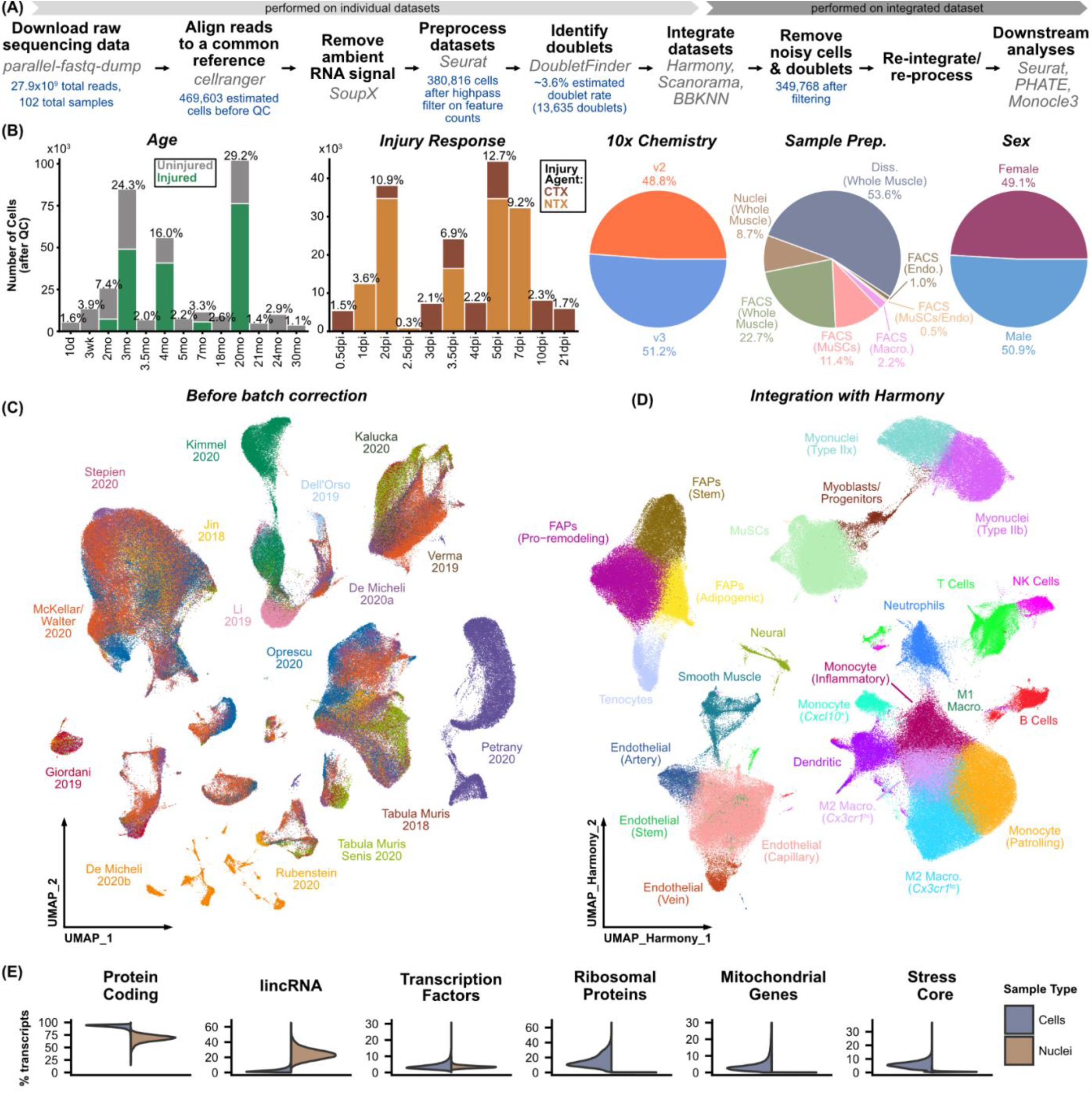
Large-scale integration of 102 scRNAseq and snRNAseq samples reveals cell subtypes in skeletal muscle. **(A)** Workflow used for preparation, integration, and analysis of sc/snRNAseq compendium (see *Methods*). **(B)** Overview of experimental and technical variables across compendium. Percentages shown are calculated with respect to cell number after quality control. Ages in months (mo). Injury by cardiotoxin (CTX) or notexin (NTX). Time-points in days post-injury (dpi). See also Table S1. **(C)** UMAP representation of the merged datasets, before batch-correction, colored by the dataset source. **(D)** UMAP representation of integrated compendium after quality filtering and batch-correction (Harmony). Cells are colored by cell type, identified after Harmony integration (Fig. S3). **(E)** Differential detection of gene biotype sets between single-cell and single-nucleus datasets, including all protein coding genes, long intergenic non-coding RNAs (lincRNAs), transcription factors, ribosomal protein subunits, mitochondrial genes, and “core” dissociation-associated stress factors.

We downloaded raw sequencing data and process them using a common pipeline [see **Methods**]. First, we re-aligned reads to a single reference genome (mm10). After removal of ambient RNA signatures (SoupX^30^), filtering of low quality cells, and identification of doublets (DoubletFinder^31^), we merged the datasets and performed initial single-cell transcriptomic analyses with Seurat^32^. Because of the range of data sources, experimental conditions, and differences in library preparation, substantial batch effects arose from differences in feature detection and read-mapping (**Fig. S1**) and resulted in dataset-specific cell populations evident in dimensionally-reduced visualizations of these data (**Fig. 1C**). To remove these batch effects, we integrated the datasets using three approaches that have been recently described-Harmony, Scanorama, and BBKNN^33–35^ (**Figs. 1D and S2)**. Integration of the datasets by Harmony revealed a cluster of cells that express canonical marker genes from multiple cell types (**Fig. S3**). Because of the enrichment of doublets and cells with high feature and transcript counts within this cluster, we reasoned that these cells were likely the result of a technical artifact and we removed them from our analyses. After removal of ambient RNA signal, doublets, and noisy cells, the resulting dataset consisted of 349,768 cells and nuclei.

After data integration, we performed shared nearest neighbor (SNN) clustering and used canonical marker genes to manually identify cell types for each batch-correction output independently (**Fig. S2B**). The size of the aggregated dataset enabled identification of subtle cell type differences. For example, we were able to distinguish the blood vessel type for endothelial cells^28^, the fiber type for mature myonuclei^9–11^, and the differentiation potential for fibro-adipogenic progenitors (FAPs^36^) (**Figs. 1D, Fig. S4**). The gene panels used for cell type classification are shown in **Fig. S4**. We found that low-resolution cell type labels (lymphocytes, myeloid cells, endothelial cells, FAPs, neural cells, smooth muscle cells, and myogenic cells) were largely consistent for each batch-correction method, but cell subtype labels varied. Variation in gene expression patterns was especially strong within the monocyte and macrophage subtypes, likely reflecting the subtle differences in transcriptional activity in these highly plastic cells. We found that Harmony performed best at integrating many heterogeneous datasets, in terms of robustness of the integration, as well as speed and memory requirements. Gene expression signatures for the cell subtypes generated by Harmony integration were the most consistent with the literature (**Fig. S4**). Harmony was also the only method able to resolve the differences between single-cell and single-nucleus datasets across all cell types (**Fig. S2**). Finally, Harmony best maintained local and global structure, especially for myogenic cells, within the dimensionally-reduced space, positioning it best for downstream analyses like pseudotime trajectory inference (**Fig. 1C)**.

We examined the effects of sample preparation (single-cell or single-nuclei) and 10x Genomics Chromium chemistry (v2 or v3) on the measured transcriptomes (**Figs. 1E, S1**). We found that the Chromium v2 and v3 chemistries provided similar sensitivity and that differences between the datasets were largely driven by sequencing depth (**Fig. S1**). In contrast, the most substantial differences were observed between the single-cell and single-nuclei preparations. These differences include an increase in intronic and intergenic reads, an increase in non-coding RNA detection, and a decrease in mitochondrial and ribosomal protein transcripts for the single-nuclei, in comparison to the single-cell preparations. Consistent with a previous analysis^10^, we observed that single-cell data is enriched for genes associated with dissociation-induced stress.

### Integrated pseudotime analysis reveals the complete trajectory of native myogenic differentiation

Previous differentiation trajectory analyses of the myogenic cell lineage have elucidated transcriptional dynamics of *in vivo* muscle regeneration^15,17^ and degeneration^37^, or *in vitro* activation^23^. However, because of the relatively small size of the datasets in these studies, the reported trajectories contained gaps within the continuum of myogenesis and contain very few cells from the most short-lived states of myogenic commitment and fusion. To fill these gaps, we selected the 79,784 myogenic cells within the integrated muscle compendium to construct a more continuous, consensus landscape of myogenesis. Importantly, the size and complexity of the myogenic cells required a scalable workflow that incorporates batch-corrected values. We found that applying the recently described tool PHATE (Potential of Heat-diffusion for Affinity-based Trajectory Embedding^38^) to dimensionally-reduce Harmony values generated continuous embeddings which reflected canonical expression patterns of myogenesis (**Fig. 2A, E**).

**Figure 2.**
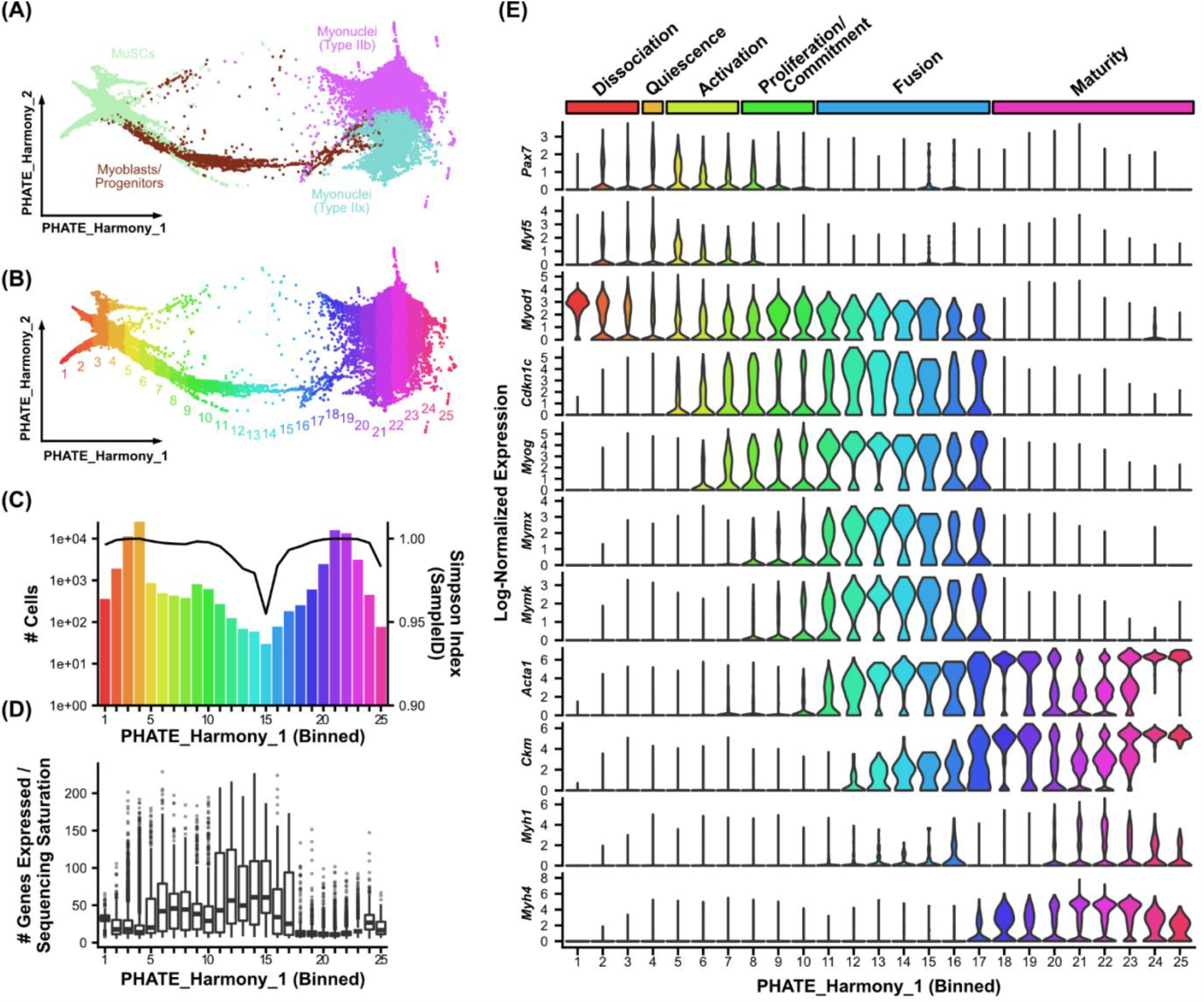
Construction of a densely sampled model of myogenesis reveals transcriptomic heterogeneity in intermediate cell states. **(A)** 79,784 myogenic and myofiber cells were selected from the integrated compendium and dimensionally reduced with PHATE to produce a consensus differentiation trajectory. **(B)** Cells were binned along the first PHATE axis (PHATE_Harmony_1). **(C)** Cell counts within each bin are shown (bar plot). Simpson’s diversity coefficients were computed on sample identifiers to determine data-source complexity for each bin (line plot). **(D)** Transcriptomic diversity by bin, reported as the number of genes detected per cell normalized to the sample sequencing saturation to account for differences in sequencing depth. **(E)** Log-normalized expression levels of canonical myogenic genes, reported as violin plots for each PHATE bin, with myogenic state detailed at top.

The PHATE embedding contained small branches emanating from the MuSCs and a distinct main branch along the first axis, toward committed progenitors and mature myofiber nuclei. Inspection of the cells in the small MuSC branches revealed that they are largely derived from samples which had been FACS-sorted and/or cultured. Consistent with previous studies showing dissociation can specifically induce stress in MuSCs^10,12,13^, these branches express apoptosis-associated genes, heat shock protein genes, and other core stress factors (**Fig. S5**). This observation suggests that dissociation-induced transcriptional responses induce distinct MuSC states that differ from native MuSC quiescence and activation, even when correcting for other technical sources of batch effect.

We next used the first PHATE dimension, derived from Harmony values, as a proxy for myogenic differentiation status (**Fig. 2B-E**). We separated the cells into 25 evenly spaced bins along the differentiation axis to visualize gene expression more easily (**Fig. 2B**). This binning demonstrates the relative sparsity of coverage and diminished data sample diversity (as reflected by a Simpson Index) within intermediate myogenic cell states, both suggesting that late fusion states are non-redundantly captured across the varied datasets within the compendium (**Fig. 2C**). Bins #1-3 contain cellular artefacts related to dissociation methods and were set aside. Canonical marker gene analysis confirmed the dataset captured cells from the major phases of myogenesis, including *Pax7*^hi^ quiescence, *Myf5*^hi^ activation, *Cdkn1c*^hi^ proliferation, *Myog*^hi^ commitment, *Mymx*^hi^ fusion, and *Acta1*^hi^ maturation (**Fig. 2D**). Consistent with previous trajectory analyses of myogenic differentiation^15,17^, the proportion of cells captured in intermediate differentiation states (defined as bins #8-17) was very small (0.7% of the compendium). Nonetheless, the large-scale integration of transcriptomic data yielded 2,623 committed or fusing cells, enabling a detailed analysis of the transcriptomic activity of these cell populations.

We next sought to explore the continual changes in myogenic differentiation and to more deeply describe MuSC commitment. Recent work has shown that the overall number of genes expressed within a cell narrows during mouse development and human mesodermal lineage specification^39^. To test whether this is also a feature of adult myogenesis, we normalized the number of features in each cell to the sequencing saturation from that sample to account for differences in sequencing depth and found that the number of genes expressed per cell is elevated in myogenic progenitor cell states (**Fig. 2D**). Notably, this burst in transcript diversity marks the exit from quiescence, continues through differentiation, and then is suppressed in mature myofibers. We found that activated (bins 5-7), committed (bins 8-10), and fusing (bins 11-17) myogenic cells were enriched within injured samples, most notably at time points beyond four days post-injury (**Fig. 3A**). Within the committed and fusing compartments (2,623 total cells), 82 of the 102 samples were represented, with between one and 131 cells captured from each sample (**Fig. 3B**). This observation again underscores the paucity of these cells within each individual sample and the power of large-scale integration for analyzing rare cell states.

**Figure 3.**
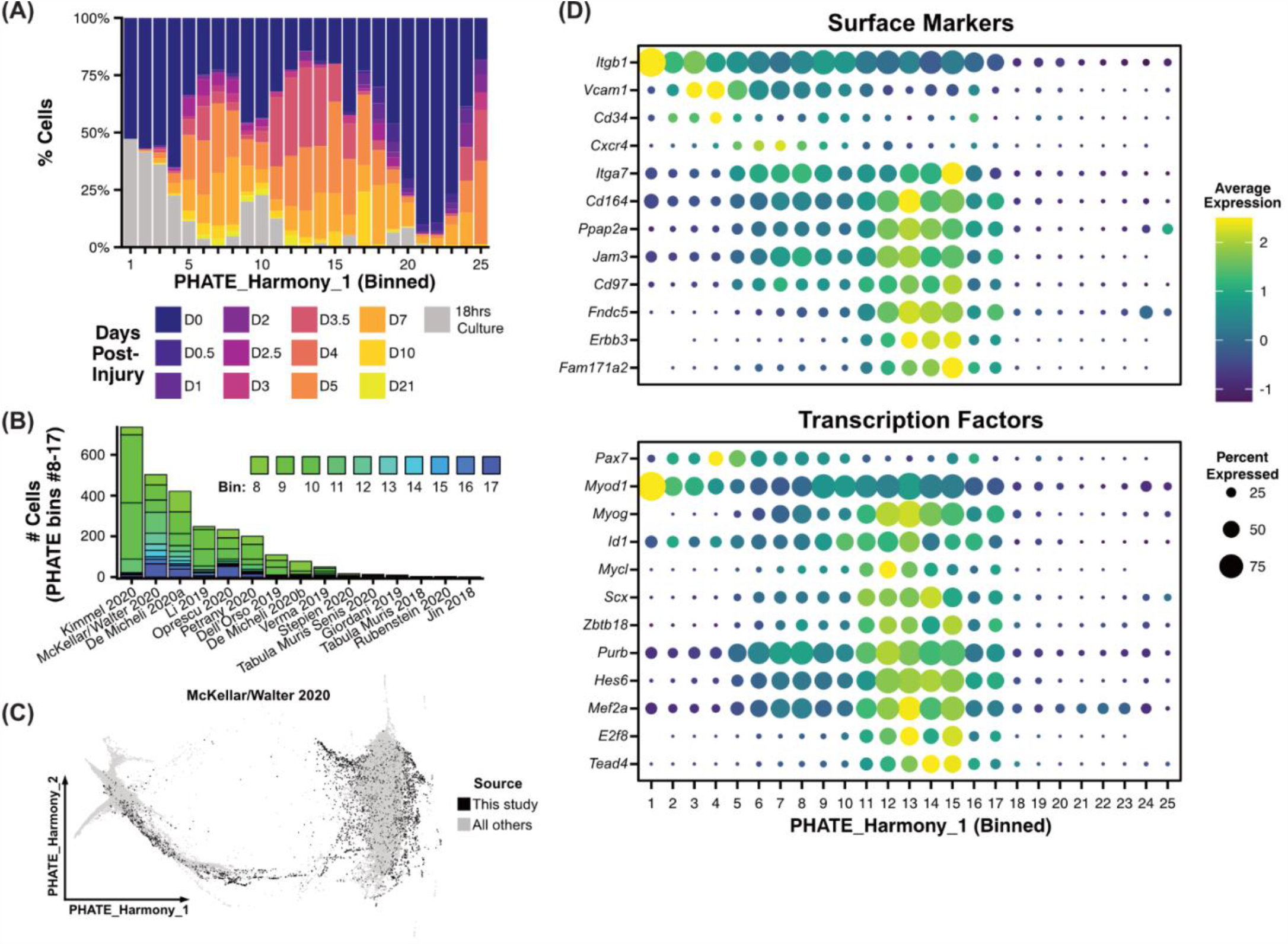
Identification of sparsely transcribed genes marking transient states in myogenesis. **(A)** Cellular composition of each myogenic PHATE bin (see Fig. 2B-C) colored by the sample’s injury time-point. **(B)** The subset of committed and fusing cells (bins 8-17; see Fig. 2E) were examined. The number of these cells in each dataset source is plotted as a stacked bar, segmented by PHATE bins (see Fig. 2E). The inset shows the PHATE embedding with the 6,648 myogenic cells newly generated in this study. **(C)** Dot plots showing the expression frequency and average level of surface markers (*top*) and transcription factors (*bottom*) genes in each PHATE bin. Canonical (*Itgb1, Vcam1, Cd34, Cxcr4, Itga7, Pax7, Myod1, Myog*) and select differentially expressed genes are reported.

Two classes of genes, surface proteins and transcription factors, have been vital in the study of MuSCs and myogenesis^3,4,40^. Surface proteins are used to isolate MuSCs and determine their regenerative potency, and the ordered expression of transcription factors has been used to define the progression of differentiation. Because these genes tend to be lowly expressed^40^, we postulated that our large-scale integrated dataset, which continually spans myogenesis, may be able to identify new markers, especially for transient committed cells. We performed differential gene expression (Wilcoxon Rank Sum test, Methods) between each of the bins along the first PHATE dimension. After filtering (adjusted p-value<10^−10^, average log_2_-fold-change>0.5), we found 2,481 genes which are differentially expressed across differentiation, including those genes associated with dissociation-induced stress. We selected surface markers among genes that were enriched in committed and fusing cells (bins 8-17), by cross-comparing against a list of surface proteins generated by the Cell Surface Protein Atlas^41^. Of the 84 resulting genes, *Cd164, Ppap2a, Jam3, Cd97, Fndc5, Erbb3*, and *Fam171a2* were specifically and highly expressed in committed and fusing myogenic cells (**Fig. 3C, Sup. File 3**). Notably, we also saw latent expression of surface markers widely used to select MuSCs in committed and fusing cells (*Itgb1, Vcam1, Cd34, Cxcr4*, and *Itga7*).

We next filtered the 2,481 differentially expressed genes for transcription factors (MGI GO term “DNA-binding transcription factor activity”) that were enriched in committed and fusing cells. Among the 100 resulting genes, we found 10 transcription factors which were highly expressed only during the commitment and fusion stages of myogenesis (*Myog, Id1, Mycl, Scx, Zbtb18, Purb, Hes6, Mef2a, E2f8, Tead4*). Overall, these findings demonstrate that large-scale integration of single-cell transcriptomic data can enable detailed gene expression profiling of rare cell states, even for lowly expressed transcripts. The surface markers identified here may be used to deplete more committed cells in MuSC isolation protocols or more selectively enrich transient committed progenitor cells. The transcription factors represent potential targets for inhibiting commitment in MuSC expansion protocols.

### Deconvolution of spatial RNA sequencing data using a large-scale, integrated reference

The biggest limitation of droplet-based scRNAseq methods is their inability to recover spatial information. Current methods for spatial RNA sequencing are limited either by the number of features they can detect or the spatial resolution which they can resolve^42^. We used the compendium to deconvolve low-resolution spatial RNA sequencing data and identify which cell types localize together during muscle injury response. We performed spatial RNA sequencing using the Visium platform (10x Genomics), which uses a patterned array of spots containing barcoded oligonucleotides to capture transcripts across a tissue slice. To enrich for the intermediate myogenic cell populations, we collected samples two, five, and seven days after chemical injury with notexin (**Fig. 4A**). Based on canonical gene expression analysis, we observed MuSC activation (*Myod1*) at 2 dpi followed by peaks in myogenic commitment markers (*Myog*) and fusogens (*Mymk*) at 5 dpi, within the muscle injury zone (**Fig. 4B-C**).

**Figure 4.**
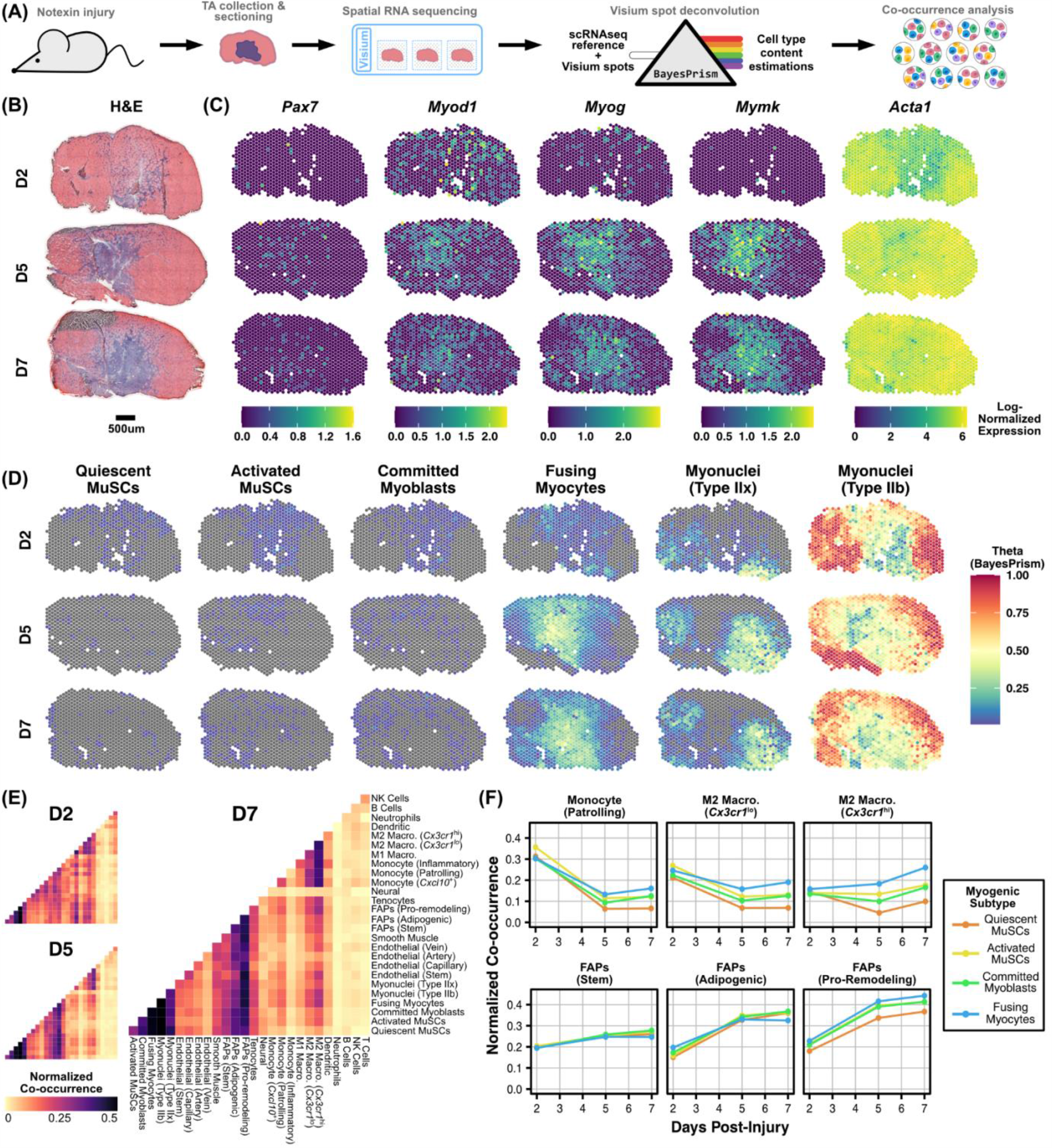
Deeply profiled cell types enable deconvolution of spatial RNA-sequencing data. **(A)** Workflow for the generation of Visium spatial RNA sequencing of regenerating skeletal muscle and spot deconvolution to cell subtype annotations from the sc/snRNA-seq compendium. **(B)** H&E images of mouse tibialis anterior samples at two, five, and seven days post-notexin injury used for spatial RNA sequencing. **(C)** Spatial expression patterns of canonical myogenesis gene expression (log-normalized) by Visium spot. **(D)** Inferred myogenic cell subtype content estimation via BayesPrism. Theta values reflect the estimated fraction of transcripts attributed to each cell type. **(E)** Spot co-occurrence between each of the 27 cell subtypes, normalized to the cumulative abundance across each row (cell subtype). Injury time-points reported separately. **(F)** Normalized spot co-occurrence of select immune and fibro/adipogenic progenitor cell subtypes with myogenic cell subtypes by injury time-point.

Each Visium spot is 55 µm in diameter and therefore contains several cells. To deconvolve each spot we used BayesPrism, a Bayesian algorithm designed to estimate cell type composition within a bulk RNAseq dataset using a single-cell reference as prior information^43^. We treated each individual spot as a bulk RNA sequencing sample and used BayesPrism to estimate what fraction of the transcripts (theta) within that spot are derived from each cell type within a single-cell reference. We defined the cell types by incorporating PHATE binning labels (**Fig. 2B**) into our single-cell reference (**Fig. 1D**). We relabeled MuSCs and Myoblasts/Progenitors (**Fig. 1D**) as Quiescent MuSCs, Activated MuSCs, Committed Myoblasts, or Fusing Myocytes, according to the PHATE bin they occupied.

Interestingly, deconvolution by BayesPrism revealed the spatial localization of subtly different cell types (**Fig. 4D**). We were able to distinguish quiescent and activated MuSCs, and other myogenic cell states. We next explored which stromal and immune populations co-localized with each myogenic cell type. We ranked the cell types in each spot by theta value, counted the top 10 cell types, and tallied the co-occurrence for each pair of cell types. We then normalized those tallies by the relative abundance of one cell type in the pair (Normalized Co-occurrence, **Fig. 4E**). We found that, at two days post-injury, “patrolling” monocytes (*Fabp5*^hi^/*Cx3cr1*^neg^/*Ly6c2*^neg^) and *Cx3cr1*^lo^ M2 macrophages (*Csf1r*^hi^/*C1qa*^hi^, also called anti-inflammatory macrophages) both co-localize with myogenic cells. For the time point later in the injury response (day 7), we observed a shift towards *Cx3cr1*^hi^ M2 macrophages co-localizing with myogenic cells, especially fusing myocytes. We also found an increase in co-localization of FAPs during the injury response, especially “pro-remodeling” FAPs (*Pdgfra*^hi^/*Cd34*^lo^/*Mmp14*^hi^/*Col3a1*^hi^). Overall, these analyses demonstrate how a large-scale single-cell reference transcriptome can be used together with spatial RNA-seq data to spatially map infrequent cell types and states.

## DISCUSSION

scRNAseq methodologies have enabled a high resolution view of cellular dynamics in muscle injury response. However, current methods are ill-suited to study rare and transient cell populations due to limitations on the number of cells captured from each sample. For the few rare cells which are captured, even deeply sequenced samples reflect only a portion of the transcriptome in each cell, because of limitations in the number of molecules captured for each cell^44^. Here, we demonstrate that large-scale integration of newly generated and public single-cell transcriptomic data can overcome these limitations. We provide a scalable pipeline of alignment, quality control, processing, and integration to unify sc/snRNAseq data from diverse experimental settings and techniques for common analysis. We annotated cell subtypes using a curated panel of 115 marker genes from the literature^11,15,16,28,36,45–55^. This approach provides a consensus resource of cell-type annotations for the skeletal muscle field, which may enable more consistent analyses across studies and allow for more clear identification of cell subpopulations. Indeed, we observed distinct subpopulations of FAPs, endothelial cells, and immune cells which were not reproducibly evident in prior individual sc/snRNAseq studies^15–17^. For example, immune cells distinctly split into neutrophils, dendritic cells, three populations of monocytes (patrolling, inflammatory, *Cxcl10*^hi^), M1 (pro-inflammatory) macrophages, two populations of M2 (anti-inflammatory) macrophages, and separate clusters of B, T, and NK cells, which are the primary immune cell constituents in muscle repair^56,57^.

We then focused our analysis on the transcriptomic changes that occur during myogenic differentiation. Though a number of prior reports have documented transcriptomic alterations between myogenic stem and progenitor cells, our integrative approach was capable of separating sample-specific artifacts from native differentiation. Furthermore, we generated a densely populated, continuous, transcriptomic model of myogenesis, including rare transitional myogenic cells. We identified candidate surface markers and transcription factor regulators distinct to the stages of myogenic commitment and myocyte fusion (represented by PHATE bins 8-10 and 11-17, respectively), which could not be resolved with individual datasets. Interestingly, we observed transcriptional signatures of surface receptors, such as *Erbb3* and *Cd97*, specific to fusing myocytes (bins 11-17), which may enable improved prospective isolation strategies compared to less stage-specific cell markers like β1-integrin (*Itgb1*). Notably ERBB3/HER3 (encoded by *Errb3*) has been identified as a myogenic progenitor marker of human pluripotent stem cell-derived myogenic progenitors^58^. Similarly, we observed a set of transcription factors with specific expression at the commitment/fusion stages, including *E2f8, Tead4*, and *Mef2a. Mef2a* has been reported to be a myogenic commitment regulator^37,59^. Intriguingly, Tead4 is required for myoblast differentiation and binds the E2F8 gene, suggesting that E2F8 may help mediate Tead4-promoted myogenic commitment^60,61^. More generally, we found that myogenic progression is characterized by a wave of transcriptomic diversification. We observed a larger number of RNA transcripts per cell, after batch-correction and accounting for sequencing depth differences across studies, at the progenitor commitment and fusion stages than in muscle stem cells or myofibers. This agrees with the observation that stem/progenitor cell trajectories exhibit changes in transcriptional diversity across their maturation axis, but differs specific trends found in hematopoesis^39^.

Lastly, we supplemented our compendium with the first, to our knowledge, transcriptome-level spatial RNA sequencing dataset of regenerating murine skeletal muscle. We repurposed BayesPrism^43^, a recently developed algorithm for the deconvolution of bulk RNA sequencing datasets, to estimate the cell composition of each spot. We then identified putative cellular interactions within those spots that may help drive myogenesis. Our findings suggest that dynamic alterations in immune-myogenic cell co-localizations may influence muscle repair outcomes. For example, the *Cx3cr1*^hi^ M2 (anti-inflammatory) macrophage subset shifts in its colocalization tendency as muscle repair proceeds, and, at 7 days post-injury, preferentially colocalizes with fusing myocytes rather than quiescent MuSCs.

The past decade has shown rapid growth in the number of cells that can be assayed in each experiment. Recent methods which utilize combinatorial indexing have generated datasets with more than 10^6^ cells at once^62,63^. Unfortunately, these methods are technically difficult, produce relatively low depth of coverage for each cell, and require huge amounts of sequencing. These factors lead to large costs which are often outside the budget of individual academic laboratories. Large collaborative efforts such as the Tabula Muris Senis or Human Biomolecular Atlas Program have generated massive reference transcriptome datasets, but have done so only for a select number of tissues and disease settings^26,27,64^. The popularization of platforms like the 10x Chromium have led to a wealth of data for many biological systems outside the purview of these consortia. We propose large-scale integration as the most economic and effective method for generating consortium-level reference transcriptomes. To enable access and interrogation of this unified transcriptomic compendium, we generated a public web tool (scmuscle.bme.cornell.edu) for explore these diverse transcriptomic data by cell populations and within the highly resolved myogenic continuum.

Large-scale integration enables the incorporation of significant sample diversity into reference transcriptomes, which will likely better reflect the underlying biology across individuals. The necessity of including diversity in reference genomes has been realized^65–67^. Should transcriptomics move from the bench to the bedside, computational tools which can include the ethnic and genetic backgrounds of the entire patient population will be critical. We have shown that batch-correction algorithms for scRNAseq can account for differences in sex, age, and sample material. We found that the increased scale of the data allows improved resolution into the underlying biology. We have also demonstrated how transcriptomic references can be used to enhance other data types, like spatial RNA sequencing, which may have either lower cellular resolution or depth of coverage. Taken together, these results suggest that a large-scale integration of single-cell transcriptomic data can increase the long-term impact of individual experiments and can be used a tool for biological discovery.

## METHODS

### Mice

The Cornell University Institutional Animal Care and Use Committee (IACUC) approved all animal protocols, and experiments were performed in compliance with its institutional guidelines. Adult C57BL/6J mice were obtained from Jackson Laboratories (#000664, Bar Harbor, ME) and were used at 4-7 months of age. Aged C57BL/6J mice were obtained from the National Institute of Aging (NIA) Rodent Aging Colony and were used at 20 months of age. For new scRNAseq experiments, female mice were used in each experiment.

### Mouse injuries and single-cell isolation

To induce muscle injury, both tibialis anterior (TA) muscles of old (20 months) C57BL/6J mice were injected with 10 µl of notexin (10 µg/ml; Latoxan, France). At 0, 1, 2, 3.5, 5, or 7 days post-injury (dpi), mice were sacrificed and TA muscles were collected and processes independently to generate single-cell suspensions. Muscles were digested with 8 mg/ml Collagenase D (Roche, Basel, Switzerland) and 10 U/ml Dispase II (Roche, Basel, Switzerland), followed by manual dissociation to generate cell suspensions. Cell suspensions were filtered through 100 and 40 μm filters (Corning Cellgro # 431752 and #431750) to remove myofiber debris. Erythrocytes were removed through incubation in erythrocyte lysis buffer (IBI Scientific # 89135-030).

### Single-cell RNA-sequencing library preparation

After digestion, single-cell suspensions were washed and resuspended in 0.04% BSA in PBS at a concentration of 10^6^ cells/ml. Cells were counted manually with a hemocytometer to determine their concentration. Single-cell RNA-sequencing libraries were prepared using the Chromium Single Cell 3’ reagent kit v3 (10x Genomics, Pleasanton, CA) following the manufacturer’s protocol. Cells were diluted into the Chromium Single Cell A Chip to yield a recovery of 6,000 single-cell transcriptomes. After preparation, libraries were sequenced on the NextSeq 500 (Illumina, San Diego, CA).

### Spatial RNA sequencing library preparation

Tibialis anterior muscles of adult (5 mo) C57BL6/J mice were injected with 10µl notexin (10 µg/ml) at 2, 5, and 7 days prior to collection. Upon collection, tibialis anterior muscles were isolated, embedded in OCT, and frozen fresh in liquid nitrogen. Spatially tagged cDNA libraries were built using the Visium Spatial Gene Expression 3’ Library Construction v1 Kit (10x Genomics, Pleasanton, CA). Optimal tissue permeabilization time for 10 µm thick sections was found to be 15 minutes using the 10x Genomics Visium Tissue Optimization Kit. H&E stained tissue sections were imaged using Zeiss PALM MicroBeam laser capture microdissection system and the images were stitched and processed using Fiji ImageJ software. cDNA libraries were sequenced on an Illumina NextSeq 500 using 150 cycle high output kits (Read 1=28bp, Read 2=120bp, Index 1=10bp, and Index 2=10bp). Frames around the capture area on the Visium slide were aligned manually and spots covering the tissue were selected using Loop Browser v4.0.0 software (10x Genomics). Sequencing data was then aligned to the mouse reference genome (mm10) using the `spaceranger` v1.0.0 pipeline to generate a feature-by-spot-barcode expression matrix (10x Genomics).

### Data and code availability

Single-nucleus RNA sequencing data were kindly provided by the Millay lab^11^, prior to public release. Newly collected scRNAseq data for 2 samples from 7 mo mice have been deposited in GEO under accession GSE159500. The data from 20 mo mice will be released in GEO upon final publication. Spatial RNA sequencing data was deposited under GSE161318. A complete list of GEO accession numbers for the new and previously published scRNAseq data can be found in **Table S1**. SRR numbers for downloading each sample are compiled in **Sup. File 1**. All code for processing and analysis of the scRNAseq and spatial RNA sequencing data, as well as supplemental data are available on Github (github.com/mckellardw/scMuscle). The full integrated data with visualization tools is available at scmuscle.bme.cornell.edu.

### Download and alignment of single-cell RNA sequencing data

For all samples available via SRA, `parallel-fastq-dump` (github.com/rvalieris/parallel-fastq-dump) was used to download raw .fastq files. Samples which were only available as .bam files were converted to .fastq format using `bamtofastq` from 10x Genomics (github.com/10XGenomics/bamtofastq). Raw reads were aligned to the mm10 reference using `cellranger` (v3.1.0).

### Preprocessing and batch correction of single-cell RNA sequencing datasets

First, ambient RNA signal was removed using the default `SoupX` (v1.4.5) workflow (`autoEstCounts` and `adjustCounts`; github.com/constantAmateur/SoupX). Samples were then preprocessed using the standard `Seurat` (v3.2.1) workflow (`NormalizeData`, `ScaleData`, `FindVariableFeatures`, `RunPCA`, `FindNeighbors`, `FindClusters`, and `RunUMAP`; github.com/satijalab/seurat). Cells with fewer than 500 features, fewer than 1000 transcripts, or more than 30% of unique transcripts derived from mitochondrial genes were removed. After preprocessing, `DoubletFinder` (v2.0) was used to identify putative doublets in each dataset, individually. BC_mvn_ optimization was used for P_K_ parameterization. Estimated doublet rates were computed by fitting the total number of cells after quality filtering to a linear regression of the expected doublet rates published in the 10x Chromium handbook. Estimated homotypic doublet rates were also accounted for using the `modelHomotypic` function. The default P_N_ value (0.25) was used. Putative doublets were then removed from each individual dataset. After preprocessing and quality filtering, we merged the datasets and performed batch-correction with three tools, independently-`Harmony` (github.com/immunogenomics/harmony) (v1.0), `Scanorama` (github.com/brianhie/scanorama) (v1.3), and `BBKNN` (github.com/Teichlab/bbknn) (v1.3.12). We then used `Seurat` to process the integrated data. After initial integration, we removed the noisy cluster (**Fig. S4**) and re-integrated the data using each of the three batch-correction tools. After a second round of integration, we found that `Harmony` still yielded a cluster of noisy cells, but `Scanorama` and `BBKNN` did not.

### Cell type annotation

Cell types were determined for each integration method independently. For `Harmony` and `Scanorama`, dimensions accounting for 95% of the total variance were used to generate SNN graphs (`Seurat`, `FindNeighbors`). SNN clustering was then performed on the output graphs (including the corrected graph output by BBKNN) using Seurat (`FindClusters`). A clustering resolution of 1.2 was used for `BBKNN` (30 initial clusters) and `Harmony` (29 initial clusters), while `Scanorama` (38 initial clusters) required a lower resolution of 0.8. Cell types were determined based on expression of canonical genes (**Fig. S4**). Clusters which had similar canonical marker gene expression patterns, but were derived from different samples, were merged.

### Pseudotime workflow

Myogenic cells were initially subset based on the consensus cell types as determined by SNN clustering on all three integration methods. Harmony embedding values from the dimensions accounting for 95% of the total variance were used for dimensional reduction with PHATE, using `phateR` (v1.0.4) (github.com/KrishnaswamyLab/phateR).

### Deconvolution of spatial RNA sequencing spots

Spot deconvolution was performed using the deconvolution module in `BayesPrism` (previously known as “Tumor microEnvironment Deconvolution”, `TED`, v1.0; github.com/Danko-Lab/TED). First, myogenic cells were re-labeled, according to binning along the first PHATE dimension, as “Quiescent MuSCs” (bin 4), “Activated MuSCs” (bins 5-7), “Committed Myoblasts” (bins 8-10), and “Fusing Myoctes” (bins 11-17). Dissociation-associated muscle stem cells were ignored and myonuclei labels were retained from as “Myonuclei (Type IIb)” and “Myonuclei Type IIx”. Next, highly and differentially expressed genes across the 27 groups of cells were identified with differential gene expression analysis using `Seurat` (`FindAllMarkers`, using Wilcoxon Rank Sum Test; results in **Sup. File 2**). The resulting genes were filtered based on average log_2_-fold change (avg_logFC > 1) and the percentage of cells within the cluster which express each gene (pct.expressed > 0.5), yielding 1,051 genes. Mitochondrial and ribosomal protein genes were also removed from this list, in line with recommendations in the `BayesPrism` vignette. For each of the cell types, mean raw counts were calculated across the 1,051 genes to generate a gene expression profile for `BayesPrism`. Raw counts for each spot were then passed to the `run.Ted` function, using the “GEP” option for input.type and default parameters for the remaining inputs. Final Gibbs theta values were used as estimates for the fraction of transcripts from each spot that were derived from each of the 27 cell types.

## ACKNOWLEDGMENTS

We thank Peter Schweitzer and colleagues in the Cornell Biotechnology Resource Center for their help with preparing the Chromium and Visium datasets. We thank the Cornell Center for Animal Resources and Education for animal housing and care. We thank Andrea De Micheli, Ern Hwei Hannah Fong, and Alexandra Dalaya for helping with mouse procedures and generating single-cell RNA sequencing datasets. We thank all investigators who made their datasets public, especially Douglas Millay for providing the single-nucleus RNA sequencing data before its publication and helpful discussions. We thank Alexandre P. Cheng, Benjamin Grodner, Hao Shi, Charles Heinke, Emily Laurilliard, Umji Lee, Tinyi Chu, and Charles Danko for helpful discussions and feedback. This work was supported by the US National Institute of Health (NIH) grant 1DP2AI138242 to IDV, and NIH grant R01AG058630 to BDC and IDV.

## AUTHOR CONTRIBUTIONS

DWM, IDV, and BDC designed the study. DWM and LDW carried out the experiments. DWM, MM, MFZW, and LDW analyzed the data. DWM and LTS designed the web resource. DWM, IDV, and BDC wrote the manuscript. All authors provided feedback and comments.

## SUPPLEMENTAL MATERIAL

**Table S1.**
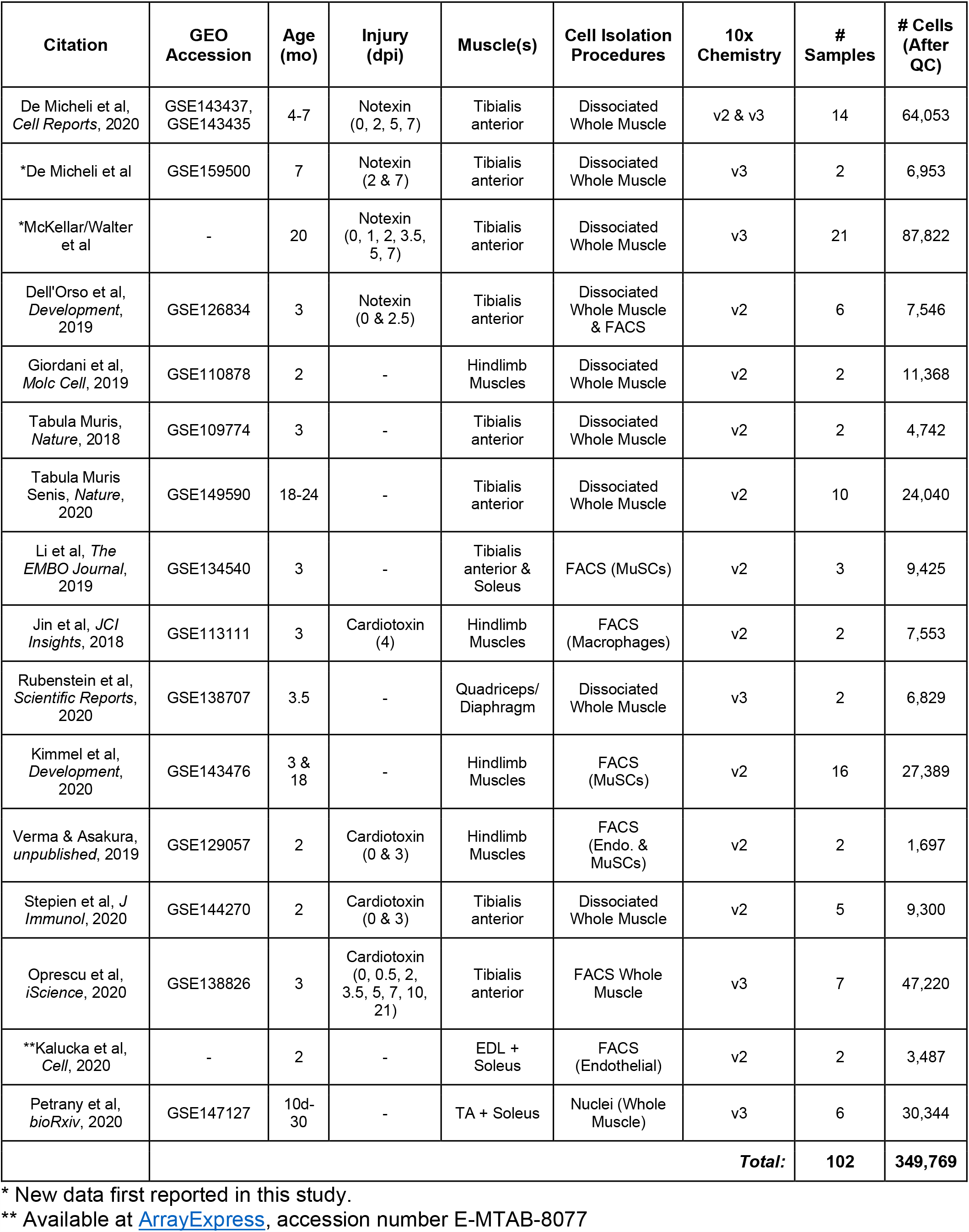
Metadata for datasets included in this study.

**Figure S1:**
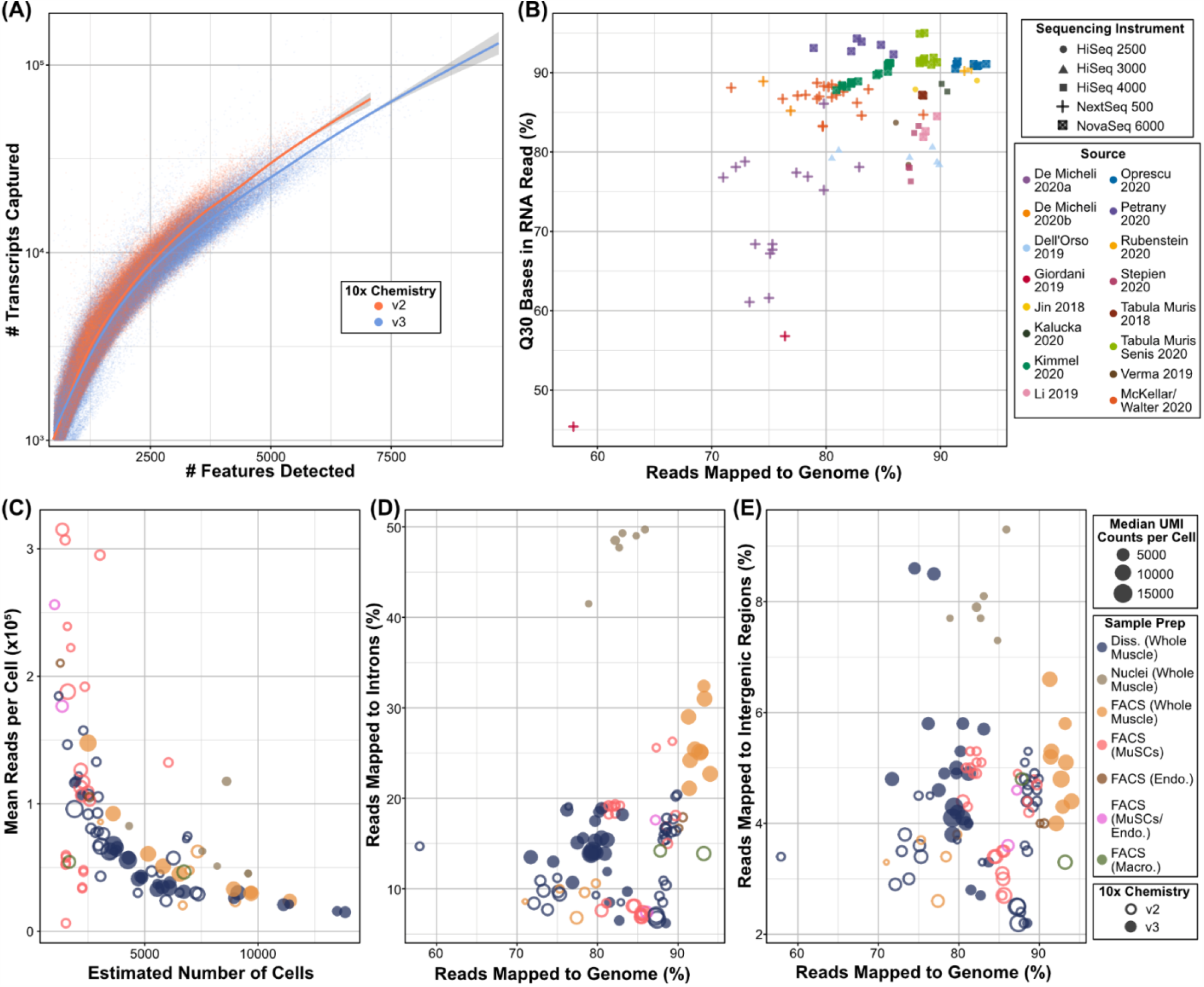
Sequencing metrics and sample material drive batch effects in single-cell RNA sequencing data. **(A)** The number of features and transcripts detected in each droplet, after quality filtering, is similar between versions 2 and 3 of the 10x Chromium chemistry. **(B)** Quality of sequencing, as determined by mappability of the library and Q30 bases inside the RNA portion of the read, may also drive batch effects. **(C)** Sequencing depth is often sacrificed in favor of higher cell counts. Single-nuclei data are enriched with **(D)** nascent transcripts and **(E)** reads outside of annotated genes.

**Figure S2.**
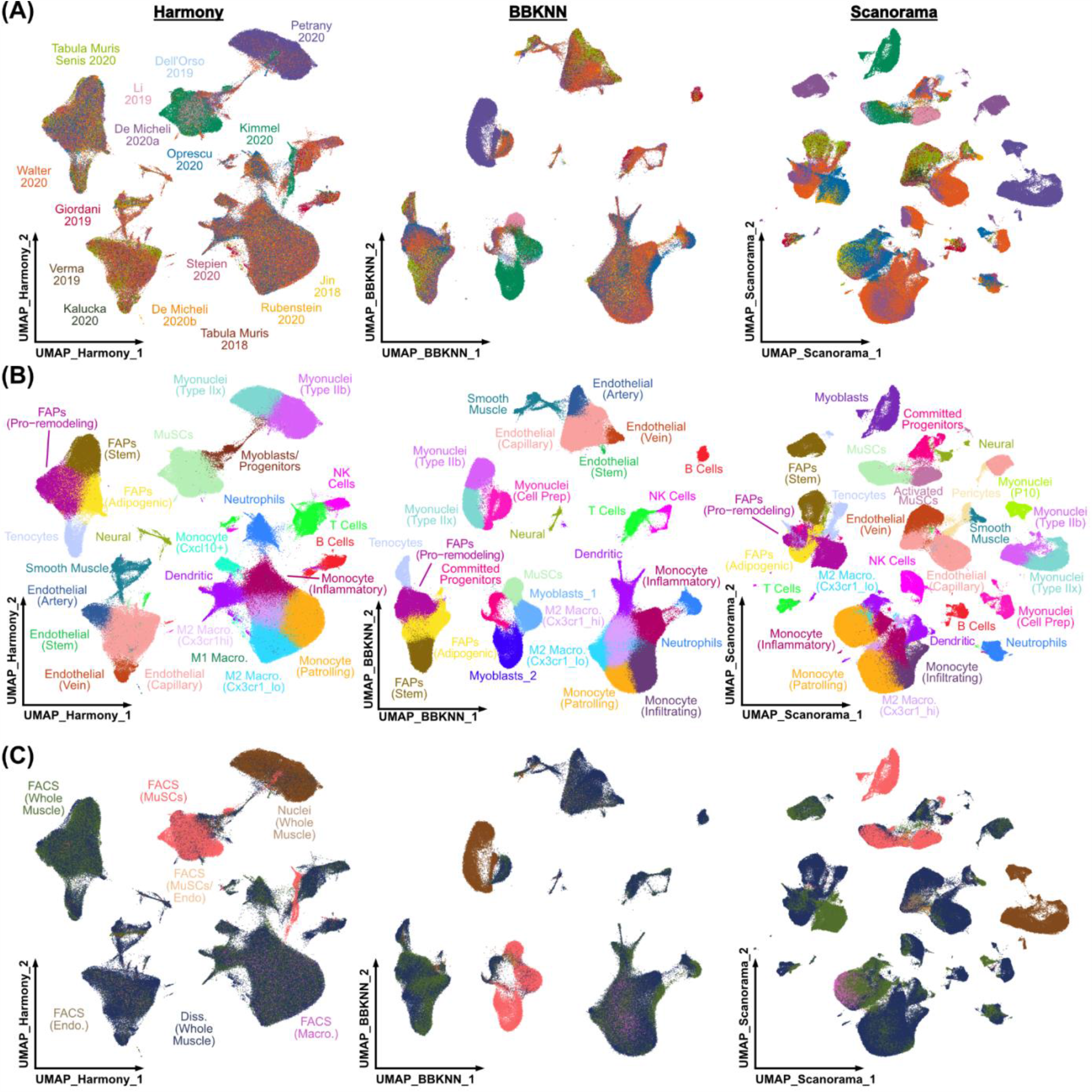
Comparison of batch-correction methods. UMAP plots generated after integration with either Harmony, BBKNN, or Scanorama are shown, colored by **(A)** source, **(B)** cell type, **(C)** or sample preparation strategy. Cell types were determined independently for each method (see Methods).

**Figure S3:**
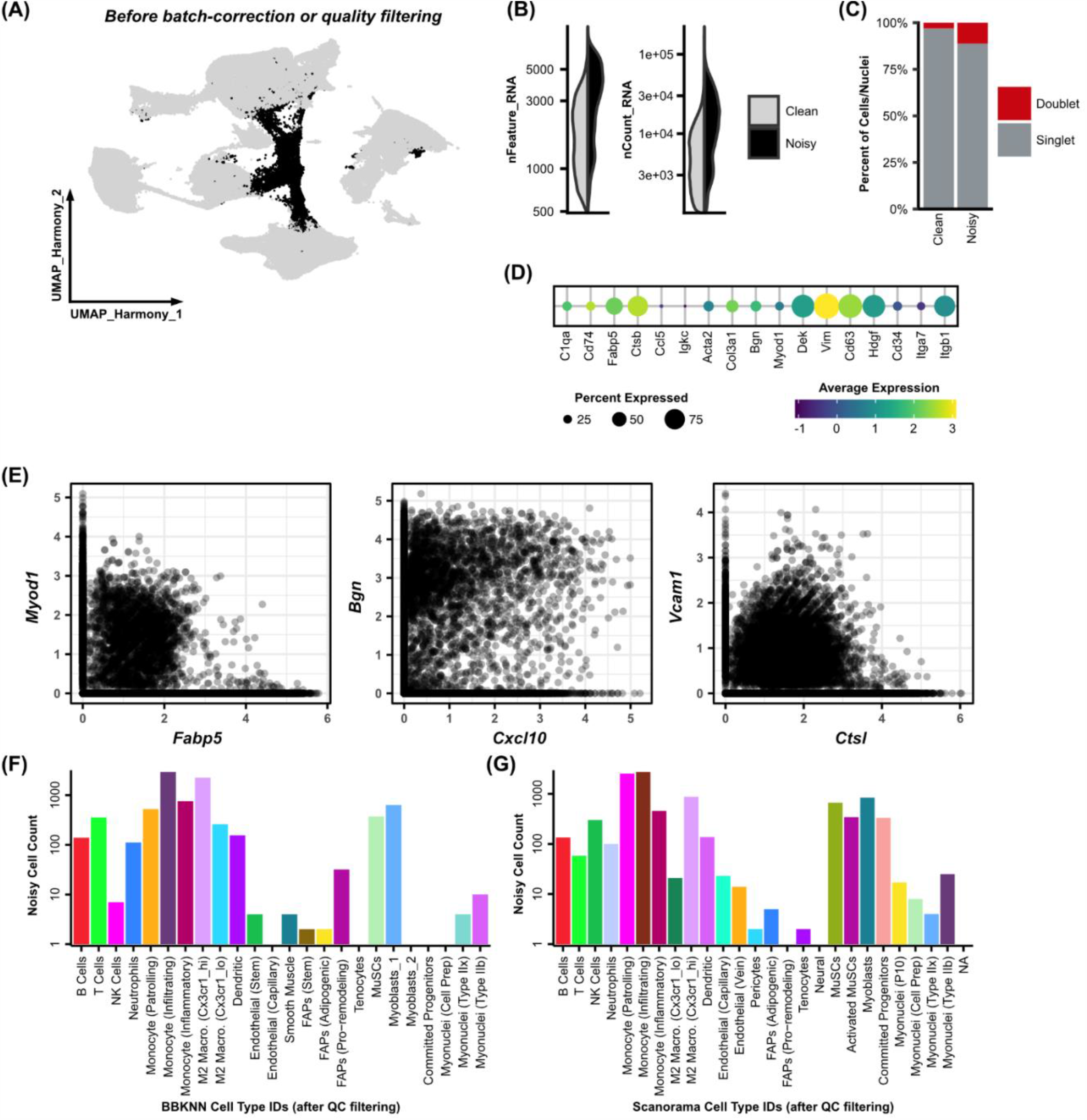
Identification of noisy cells in large-scale integration with Harmony. **(A)** After integration with Harmony and shared nearest neighbor clustering, one cluster of noisy cells with no discernible marker genes emerged. **(B)** The “Noisy” cluster is enriched with cells containing high numbers of detected features and high transcript counts. **(C)** “Noisy” cells have a four-fold enrichment for doublets, as identified by DoubletFinder, compared to other cells. **(D)** “Noisy” cells express canonical marker genes from multiple cell types. **(E)** Individual cells express multiple cell-type-specific marker genes. **(G)** “Noisy” cells which cluster together after Harmony integration are distributed across cell types after BBKNN integration and **(H)** Scanorama integration. Counts of “noisy” cells are shown as bar graphs, with a log scale, for each cluster from alternate batch-correction methods.

**Figure S4.**
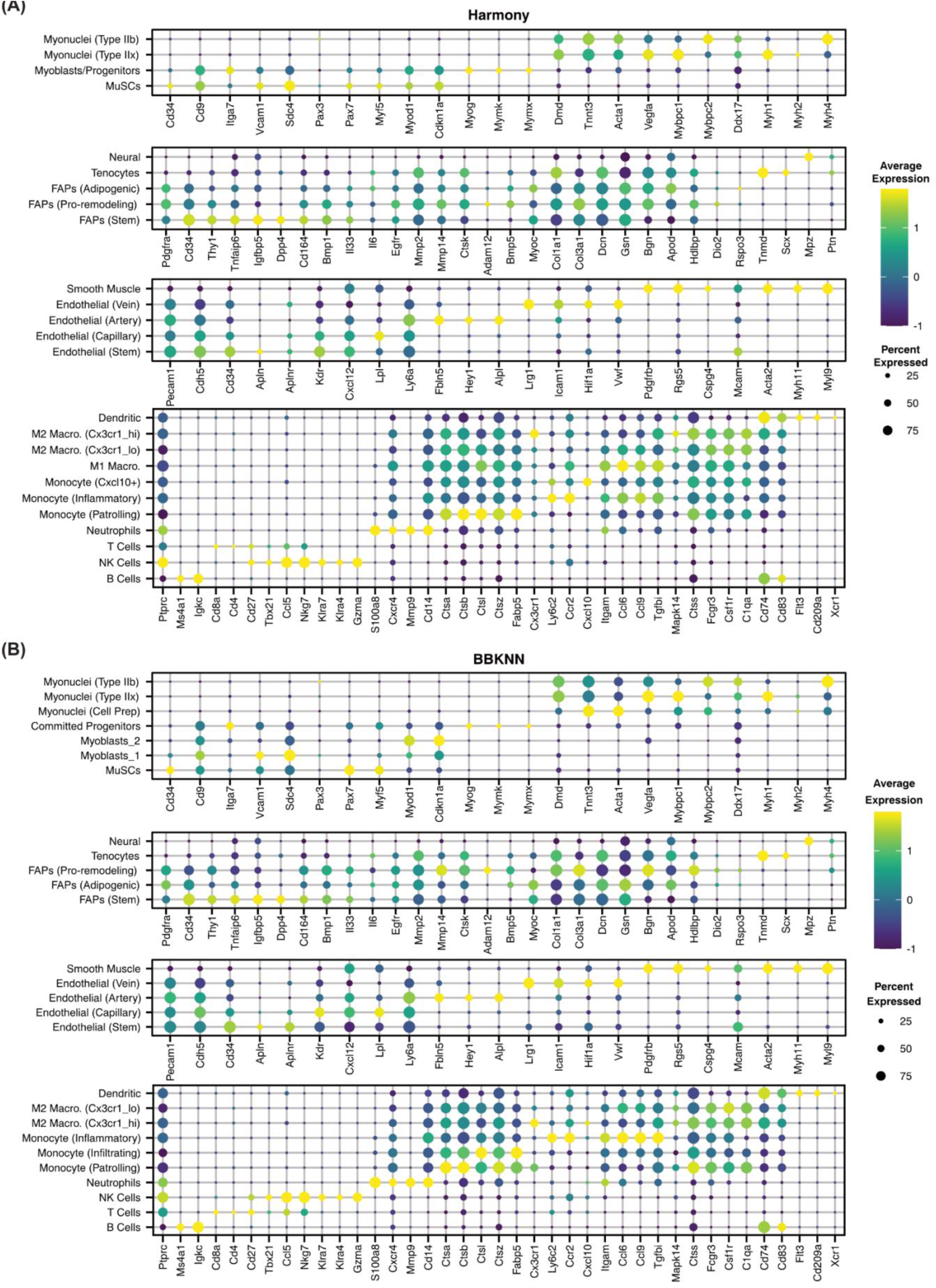
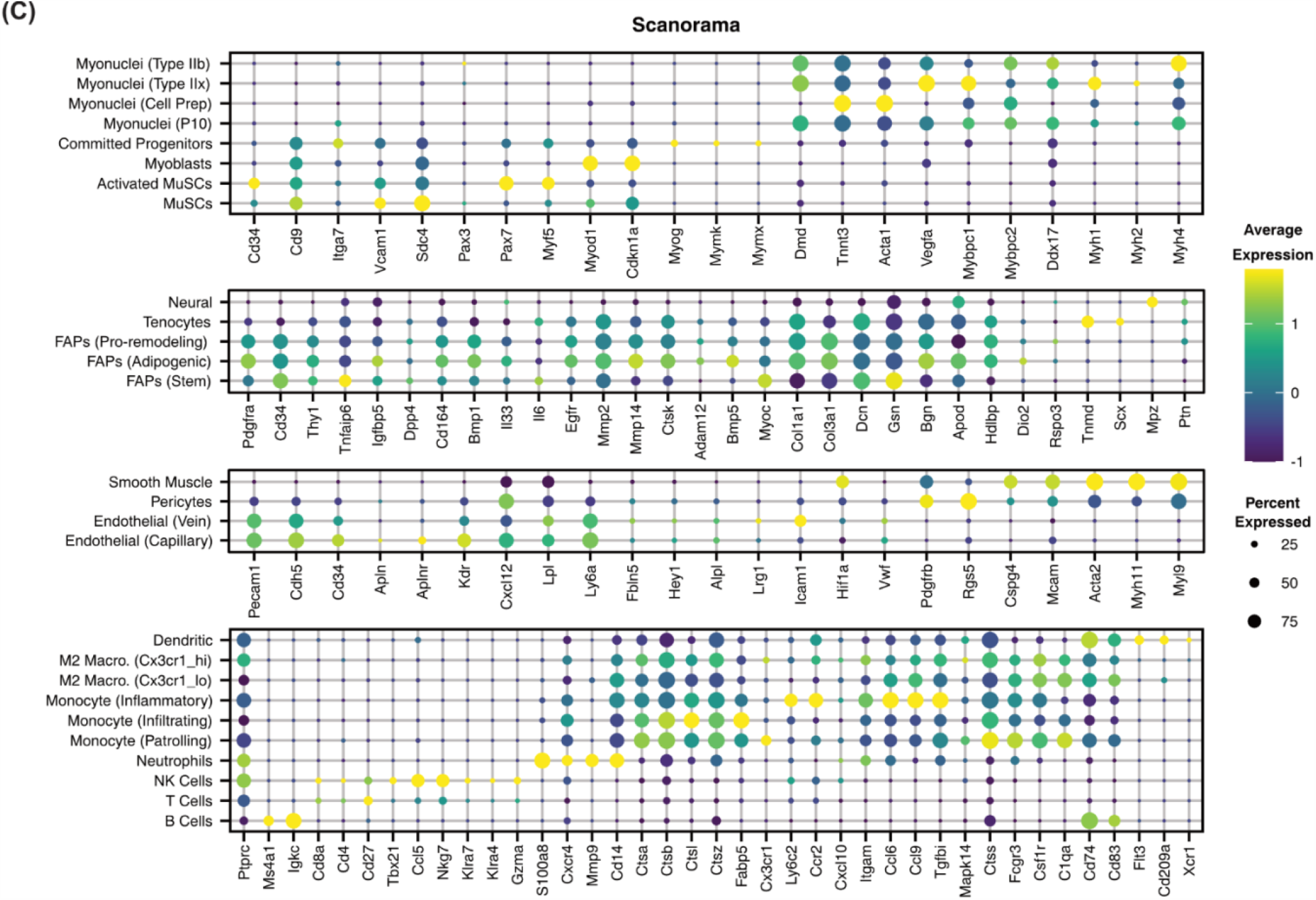
Canonical marker gene expression identifies cellular subtypes in clustering results from three batch-correction methods. Dot plots show the expression of 115 genes, curated from a literature search for cell type markers in skeletal muscle. Cell types shown were generated through shared nearest neighbor clustering on the output values from **(A)** Harmony, **(B)** BBKNN, and **(C)** Scanorama. Gene expression values are scaled (via Seurat).

**Figure S5.**
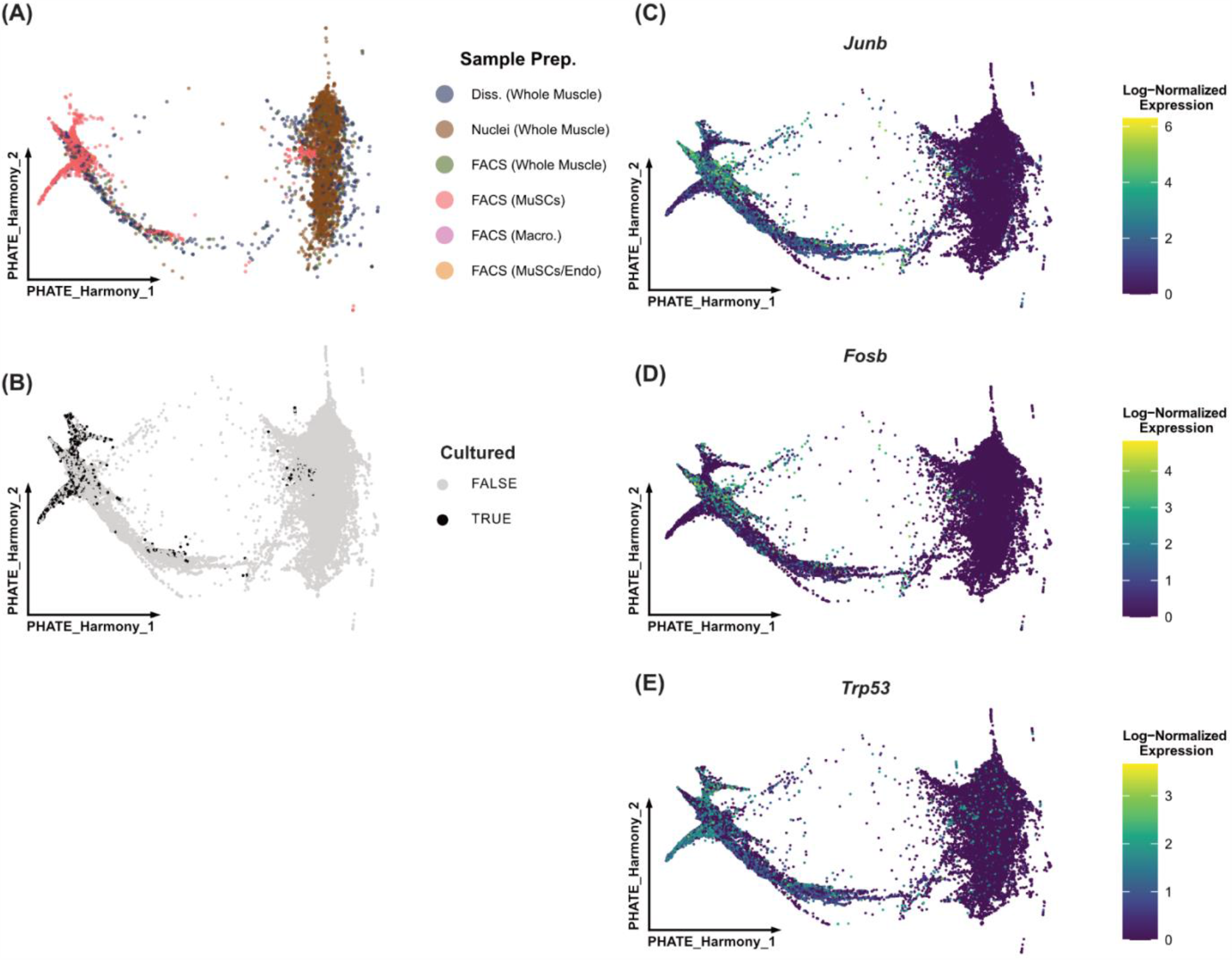
**(A)** PHATE embedding, colored by sample preparation strategy. **(B)** PHATE embedding showing which cells were FACS-sorted. **(C)** Feature plots showing expression of genes related to apoptosis (*Junb, Fosb*) and DNA repair (*Trp53*).

**Figure S6.**
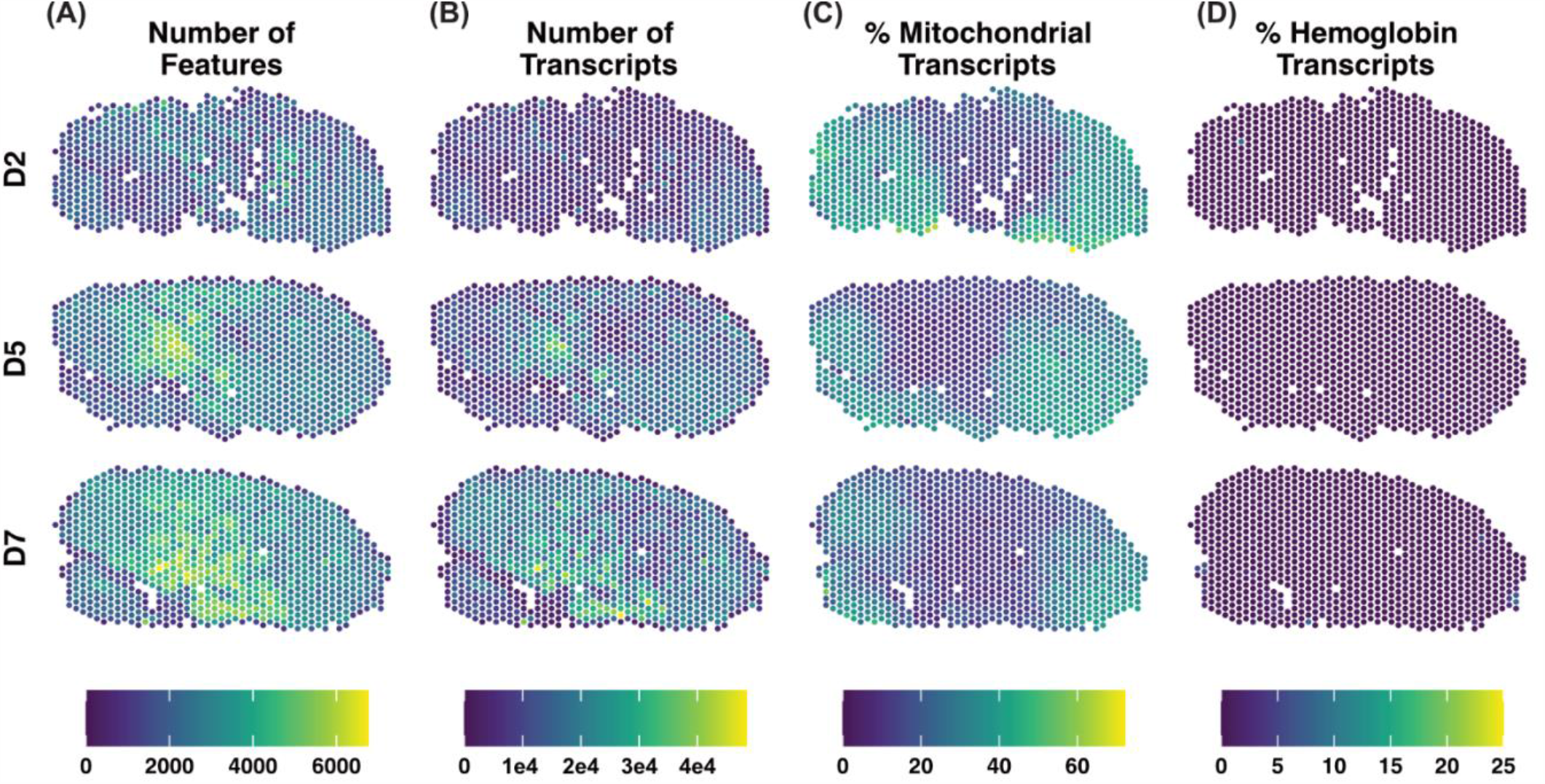
Quality metrics of Visium spatial RNA sequencing datasets. **(A)** Number of features detected, **(B**) number of transcripts detected, **(C)** percent of unique transcripts mapping to mitochondrial genes, and **(D)** percent of unique transcripts mapping to hemoglobin genes, plotted for each spot. Samples shown are tibialis anterior muscles, collected two (D2), five (D5), or seven (D7) days post-injury (notexin).

